# Mapping the spatial transcriptomic signature of the hippocampus during memory consolidation

**DOI:** 10.1101/2023.01.18.524576

**Authors:** Yann Vanrobeys, Utsav Mukherjee, Lucy Langmack, Ethan Bahl, Li-Chun Lin, Jacob J Michaelson, Ted Abel, Snehajyoti Chatterjee

## Abstract

Memory consolidation involves discrete patterns of transcriptional events in the hippocampus. Despite the emergence of single-cell transcriptomic profiling techniques, defining learning-responsive gene expression across subregions of the hippocampus has remained challenging. Here, we utilized unbiased spatial sequencing to elucidate transcriptome-wide changes in gene expression in the hippocampus following learning, enabling us to define molecular signatures unique to each hippocampal subregion. We find that each subregion of the hippocampus exhibits distinct yet overlapping transcriptomic signatures. Although the CA1 region exhibited increased expression of genes related to transcriptional regulation, the DG showed upregulation of genes associated with protein folding. We demonstrate the functional relevance of subregion-specific gene expression by genetic manipulation of a transcription factor selectively in the CA1 hippocampal subregion, leading to long-term memory deficits. This work demonstrates the power of using spatial molecular approaches to reveal transcriptional events during memory consolidation.

## Introduction

Activity-dependent gene expression occurs in wave-like patterns following experience. The early wave of transcriptional events involves increased expression of immediate early genes (IEGs) and newly synthesized proteins to regulate downstream gene expression ^1–3^. IEGs encoding transcription factors, such as *Fos, Egr1,* and the *NR4a* subfamily, regulate a larger, more diverse set of effector genes that mediate the structural and functional changes underlying synaptic plasticity. Gene expression at these critical time points is essential to drive responses to experience, including memory consolidation. Newly formed memory is thought to be stored within functionally connected neuronal populations, known as engram ensembles ^4–7^, in the hippocampal network, then gradually consolidated across multiple brain regions ^4,8–10^. Dynamic gene expression patterns represent hippocampal engram ensembles and the circuitry supporting memory consolidation ^10,11^. Neuronal populations contributing to engram ensembles are activated by learning and endure cellular changes ^10,12^, which can later be reactivated for memory retrieval ^13^ or inhibited inducing memory impairments ^14^. Therefore, understanding the transcriptional dynamics within the hippocampal circuit following an experience would provide important insights into the molecular mechanism underlying memory consolidation.

The circuitry within different subregions of the dorsal hippocampus has distinct roles in memory consolidation ^15–17^. Layer II of entorhinal cortex (EC) projects to granule cells of the dentate gyrus (DG) and pyramidal neurons of CA3 region through the perforant pathway (PP), and layer III of EC projects to the pyramidal neurons of CA1 through the temporoammonic and alvear pathways ^18–20^. The direct EC input to CA1 is essential for spatial memory consolidation and novelty detection ^21–24^. DG granule cells project onto CA3 pyramidal neurons through mossy fibers, and CA3 pyramidal neurons send projections to CA2 and CA1 pyramidal neurons through the Schaffer collateral (SC) pathway ^25–27^. The axons from CA1 pyramidal neurons project onto subiculum and EC neurons, forming the major output pathway of hippocampal circuits ^28^. The DG is the site of adult neurogenesis in the hippocampus ^29^. Adult newborn granule cells mediate pattern separation in the DG ^30^, while mature granule cells in DG and CA3 pyramidal neurons are essential for pattern completion, involving associative memory recall from a partial cue ^31,32^. Thus, hippocampal memory relies on the association between items and contexts ^33^, with neurons in the CA1 processing information about objects and locations ^34^ and DG neurons driving pattern separation to reduce overlap between neural representations of similar learning experiences ^35–37^. Despite the importance of circuitry in the dorsal hippocampus, spatial transcriptomic changes in response to learning across subregions of the dorsal hippocampus remain largely unknown.

Learning-induced gene expression has previously been shown using the whole hippocampus ^38,39^, CA1 ^40,41^, DG ^42,43^, and hippocampal neuronal nuclei ^41,44,45^, but has not been examined across all subregions simultaneously. Hippocampal engram ensembles have been studied using the expression of individual IEGs ^11^, while recent studies have applied targeted recombination of active neuronal populations to study unbiased cell-type specific gene expression in the hippocampus following a learning experience ^4,45^. *Fos* is one IEG that is thought to link hippocampal engram and place codes underlying spatial maps ^7,46^. Single nuclei RNA sequencing was recently utilized to demonstrate downstream targets of *Fos* in CA1 pyramidal cells following neuronal stimulation ^47^ and define the role of cell type-specific activity-driven expression of *Fos* in CA1 for spatial memory ^7,46,47^. Single-nuclei transcriptomic studies from Fos+ (activated) and Fos- (non-activated) hippocampal neurons following a novel environment exposure revealed transcriptomic differences between DG and CA1 neurons ^48^. Other studies have applied a similar approach in the hippocampus to capture engram cells following learning ^45^ or activated neurons following neuronal stimulation ^2,43^. However, it is still unclear how gene expression in each of the spatially and functionally distinct subregions is regulated after learning. The transcriptomic diversity within these subregions needs to be examined more clearly to better understand the role each of these subregions in memory consolidation.

Advancements in single-cell RNA sequencing analyses allows us to sort transcriptional profiles into cell types based on canonical marker genes ^49,50^. However, utilizing spatial coordinates within intact brain tissue enables precise identification of transcriptomic changes at high spatial resolution ^51,52^. Visium spatial transcriptomics (*10X Genomics*) combines both histology and spatial profiling of RNA expression to provide high-resolution transcriptomic characterization of distinct transcriptional profiles within individual brain subregions ^53^. We have recently used this Visium spatial transcriptomic approach to demonstrated neuronal activation patterns within brain regions following spatial exploration using a deep-learning computational tool ^54^. In this work, we have extended this novel approach to examine activity-driven spatial transcriptomic diversity within the hippocampal network. We define genome-wide transcriptomic changes in the CA1 pyramidal layer, CA1 stratum radiatum, CA1 stratum oriens, CA2+3 pyramidal layer, and dentate gyrus (DG) granular and molecular layers of the dorsal hippocampus within the first hour following spatial exploration. Moreover, we functionally validated our findings by selectively manipulating the function of Nr4a transcription factor subfamily members within CA1 pyramidal neurons. Mapping the precise expression patterns of genes in hippocampal subregions at an early timepoint after learning has enhanced our understanding of their role in memory consolidation.

## Results

### Pseudobulk analysis of hippocampal spatial transcriptomics following learning correlates with bulk RNA sequencing

The growing knowledge of transcriptomic heterogeneity in hippocampal subregions raises the critical question of the gene expression dynamics during a critical early timepoint of memory consolidation. To understand the learning-induced gene expression patterns exhibited by different hippocampal subregions, we performed spatial transcriptomic analyses using the *10x Genomics Visium* platform in coronal brain slices obtained from adult C57BL/6J male mice 1 hr after training in a hippocampus-dependent learning task compared to homecage controls (Spatial object recognition task, SOR, n=4/group, **Fig. 1a**). We and others have previously demonstrated that the learning-induced early wave of gene expression peaks at this timepoint after learning ^55–57^. We further examined the expression profiles by integrating our previous spatial transcriptomics dataset following SOR training ^54^ (n=3/group). We first obtained cumulative transcriptomic profiles (pseudobulk analysis, total n=7/group) by combining the hippocampal subregions CA1 pyramidal layer, CA1 stratum radiatum, CA1 stratum oriens, CA2 and CA3 pyramidal layers and DG granular layers (**Fig. 1b**). Differential gene expression analysis of this pseudobulk data revealed 101 upregulated and 18 downregulated genes following learning (**Fig. 1c-d**). Enrichment network analysis was used to identify the pathways most represented among the differentially expressed genes. The upregulated pathways include nuclear receptor activity, nucleotide transmembrane transporter activity, protein kinase inhibitor activity, dioxygenase activity and histone demethylase activity (**Fig. 1e**). The nuclear receptor activity members *Nr4a1*, *Nr4a2* and *Nr4a3* comprised a subfamily of transcription factors known to be involved in learning and memory ^58,59^. Histone demethylation activity has been linked to memory consolidation ^60^, while mutations in *Jmjd1c* are associated with intellectual disability ^61^. Protein kinase inhibitors are often found to be upregulated following learning, acting as a negative regulator of transcription activation pathways, such as MAPK pathway ^62^, and potential activation of memory suppression genes ^63^. Other immediate early genes upregulated following learning include *Egr1*, *Arc*, *Homer1*, *Per1*, *Dusp5*, and *Junb* and are all associated with learning and memory ^2,38,64^.

**Figure 1:**
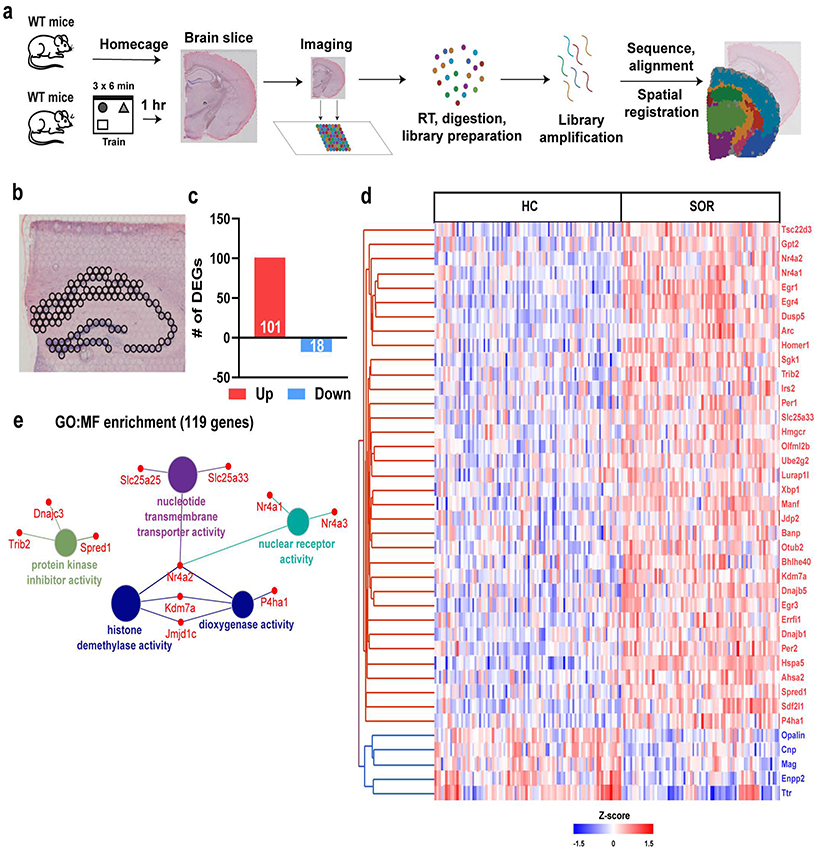
Pseudobulk RNA-seq analysis of spatial transcriptomic data defines learning-induced gene expression in the hippocampus. **a.** Schematic of the spatial learning paradigm, followed by a graphic description of the Visium pipeline. n=4/group, males only **b.** Visual depiction of spots across all the hippocampal subregions used for pseudobulk RNA-seq analysis. **c.** Bar graph illustrating the total number of upregulated and downregulated genes computed from the pseudobulk RNA-seq data. **d.** Heat map generated from individual Visium spots of the 40 top significant differentially expressed genes after learning. Red: upregulated, and blue: downregulation genes. **e.** Gene Ontology (GO) enrichment analysis performed on all the differentially expressed genes based on their molecular function (MF).

Over the past decade, bulk RNA sequencing (RNA-seq) has been extensively used to study transcriptional profiles from brain tissue ^40,41,58^. Therefore, to validate our spatial transcriptomic approach with conventional transcriptomic tools, we performed RNA-seq using whole dorsal hippocampus tissue (bulk RNA-seq) from mice trained in SOR (1 hr) or homecage. Bulk RNA-seq analysis revealed differential expression of 224 genes (DEGs, FDR<0.05) following SOR training compared to control mice, with 147 upregulated and 77 downregulated genes after learning (**Fig. 2a**). We next asked whether our pseudobulk spatial transcriptomics data overlapped with learning-induced gene expression changes observed using the bulk RNA-seq approach. Among the 101 upregulated genes from pseudobulk spatial transcriptomics, 29 genes were identified with bulk RNA-seq. Only one gene among 18 downregulated genes appeared in bulk RNA-seq. Genes differentially expressed in pseudobulk RNA-seq significantly correlate with bulk RNA-seq, and the directionality of the change in expression was maintained (**Fig. 2b**). Of these, *Nr4a1*, *Egr1*, *Egr4*, *Dusp5*, *Arc,* and *Sgk1* were among the top common upregulated genes, while oligodendrocyte differentiation-related gene *Opalin* was the only common downregulated gene (**Fig. 2b**). Pseudobulk analysis also revealed differentially expressed genes that were not identified by bulk RNA-seq approach. Some of the novel upregulated transcripts identified using pseudobulk spatial transcriptomics include genes related to chromatin binding (*Ncoa2*, *Polg*, *Smc3*, Bcl6, *Jdp2*, *Sp3*), protein kinase inhibitors activity (*Spred1*, *Trib2*) and chaperone binding (*Dnajc3*, *Sacs*, *Grpel2*). Some of the novel downregulated genes included myelin oligodendrocyte glycoprotein (*Mog*), myelin associated glycoprotein (*Mag*) and long noncoding RNA, *Mir9-3hg*. These results suggests that spatial transcriptomics using the Visium platform provides findings that overlap with other transcriptomic approaches yet reveals new genes that may be undetectable in other techniques.

**Figure 2:**
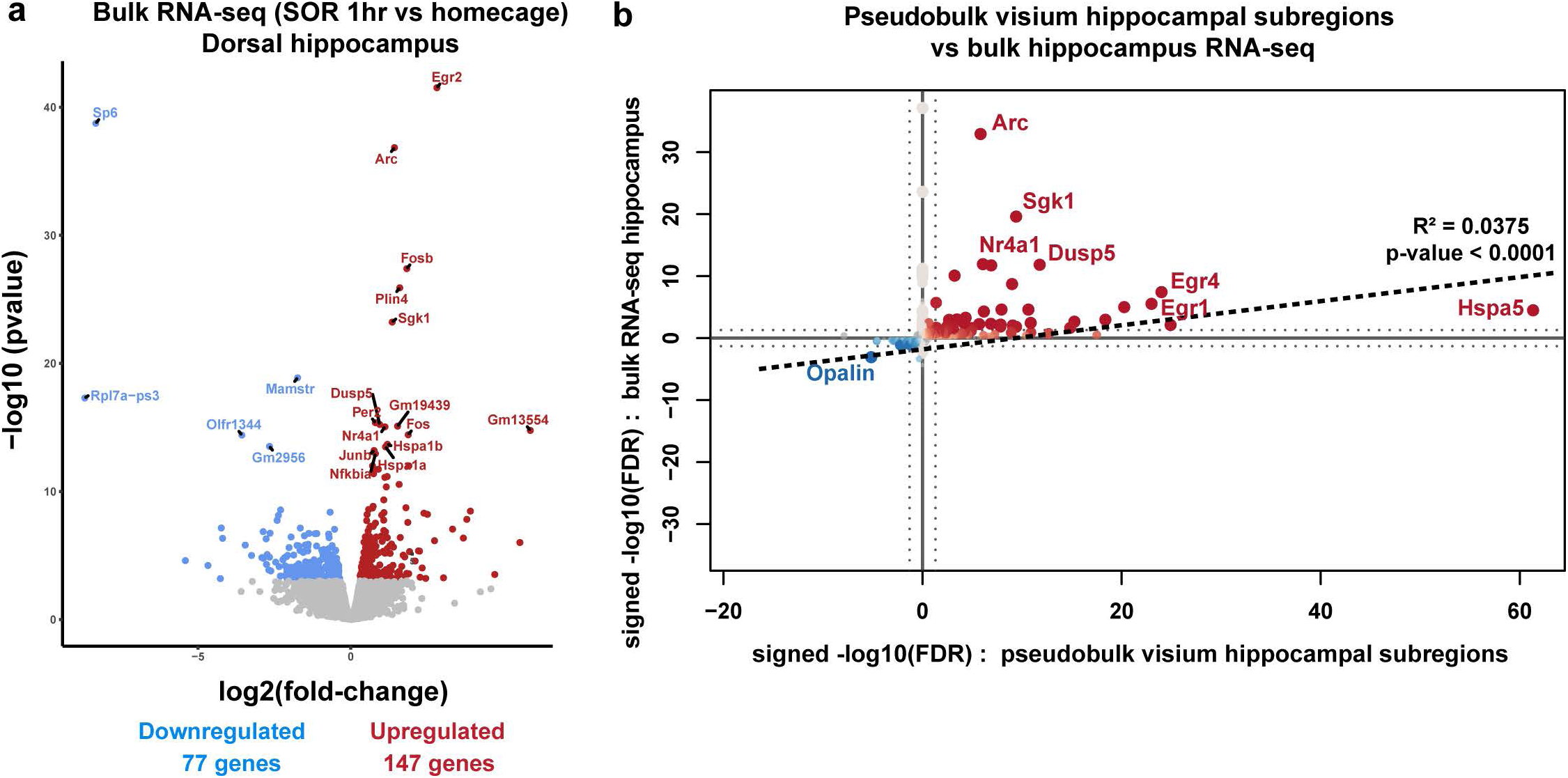
Comparison of the pseudobulk RNA-seq with the bulk RNA-seq dataset after learning. **a.** Volcano plot illustrating the most significant differentially expressed genes after learning from a bulk RNA-seq experiment performed from the dorsal hippocampus 1 hour after learning. homecage (n=4), SOR (n=4). **b.** Quadrant plot depicting the correlation between differentially expressed genes identified in bulk RNA-seq and pseudobulk RNA-seq.

### Hippocampal subregions exhibit distinct transcriptomic signatures following learning

The dorsal hippocampus is composed of multiple anatomically and functionally distinct subregions. Here we distinguished the major principal neuronal layers and memory-relevant hippocampal regions: CA1 pyramidal layer, CA1 stratum radiatum, CA1 stratum oriens, CA2 and CA3 pyramidal layers combined, and DG granular layer based on the spatial topography by H&E staining (**Fig. 3a**). Computational analysis of the transcriptomic profiles from these hippocampal subregions reveals distinct clusters in a UMAP plot (**Fig. 3b**). Analyzing the hippocampal subregion-specific transcriptomic signature after learning revealed 58 differentially expressed genes in the CA1 pyramidal layer, 16 genes in the CA2 and CA3 pyramidal layers, and 104 genes in the DG molecular and granular layer. Among these differentially expressed genes, learning induced 46 upregulated and 12 downregulated genes in the CA1 pyramidal layer, 13 upregulated and 3 downregulated genes in CA2 and CA3 pyramidal layers, and 68 upregulated and 36 downregulated genes in DG (**Fig. 3c)**. In addition to the CA1 pyramidal layer, we also investigated the transcriptomic signature exhibited by CA1 stratum radiatum and stratum oriens. CA1 stratum radiatum is the suprapyramidal region containing apical dendrites of pyramidal cells where CA3 to CA1 SC connections are located. CA1 stratum oriens is the infrapyramidal region containing basal dendrites of pyramidal cells where some CA3 to CA1 SC connections are located. However, heterogenous population of interneurons and other non-neuronal cells are also scattered through these layers. Differential gene expression analysis from these CA1 regions identified 10 upregulated and 1 downregulated gene in stratum radiatum and 9 upregulated and 9 downregulated genes in stratum oriens (**Fig. 3c)**. Enrichment network analysis revealed that the pathways enriched in the CA1 pyramidal layer include nuclear receptor activity and MAP kinase tyrosine/serine/threonine phosphatase activity (**Fig. 3d**). In contrast, the pathways in DG include protein kinase inhibitor activity and protein disulfide isomerase activity (**Fig. 3e**). Next, we utilized an upset plot to compare the differentially expressed genes from each hippocampal subregion (**Fig. 4c-d**). This analysis identified 51 genes that were exclusively upregulated in DG, 22 genes exclusively upregulated in the CA1 pyramidal layer, and 11 genes were upregulated in both CA1 and DG, but not in other hippocampal subregions (**Fig. 3f**). Some of these 11 common genes are involved in protein folding (*Xbp1*, *Sdf2l1*, *Dnajb1*) and the MAPK pathway (*Spred1)*. Genes related to activity-driven transcription regulation and MAPK pathway regulation (*Arc*, *Nr4a2*, *Per1,* and *Dusp5*) were upregulated both in CA1 and CA2+CA3 pyramidal layers, while *Nr4a1* and *Egr3* were upregulated in the CA1 pyramidal layer, stratum radiatum and stratum oriens. These findings suggest large-scale transcriptional changes in DG, while CA pyramidal region showed increased activation state of IEGs linked to engram ensemble following spatial learning.

**Figure 3:**
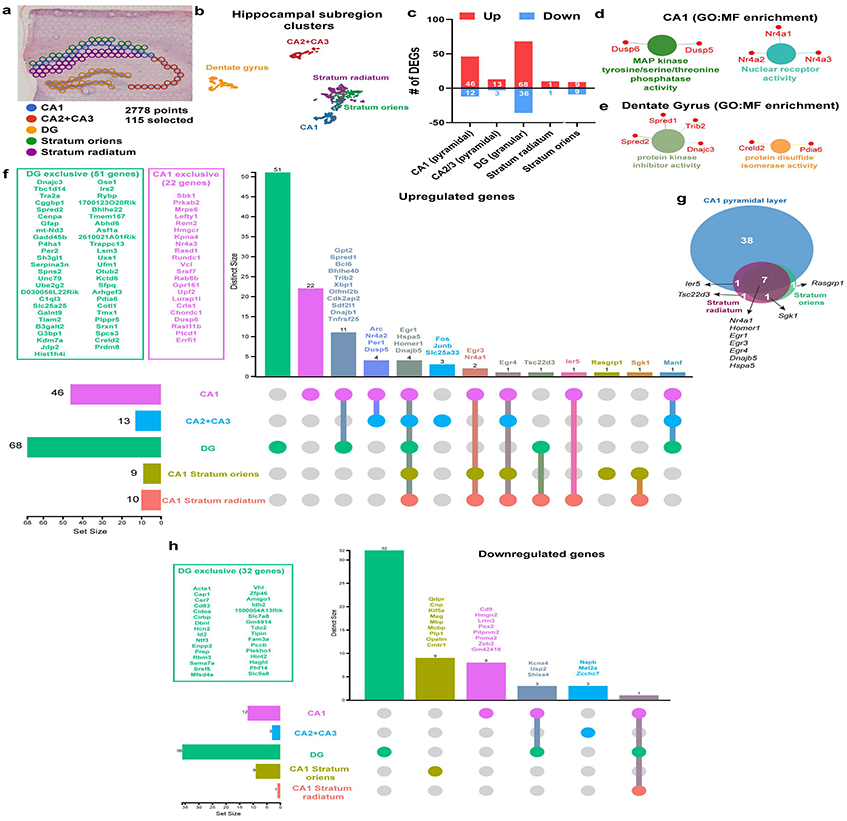
Utilizing spatial transcriptomics to dissect subregion-specific transcriptomic signature of learning in the hippocampus. **a.** Representative depiction of the Visium spots considered to distinguish hippocampal subregions. **b.** UMAP plot showing spot-clusters demarcating the most prominent hippocampal subregions. **c.** Bar graph depicting the total number of differentially expressed genes corresponding to hippocampal subregions. **d.** Gene Ontology (GO) enrichment analysis performed on the differentially upregulated genes in area CA1 pyramidal layer. **e.** Gene Ontology (GO) enrichment analysis of all differentially upregulated genes in Dentate Gyrus (DG). **f.** UpSet plot illustrating the spatial pattern of all the significantly upregulated learning-induced genes throughout the hippocampus. **g.** Venn diagram showing the overlap of upregulated genes exclusive to area CA1 pyramidal layer, Stratum Oriens, and Stratum Radiatum. **h.** UpSet plot depicting the spatial map of all the significantly downregulated genes in the hippocampus.

**Figure 4:**
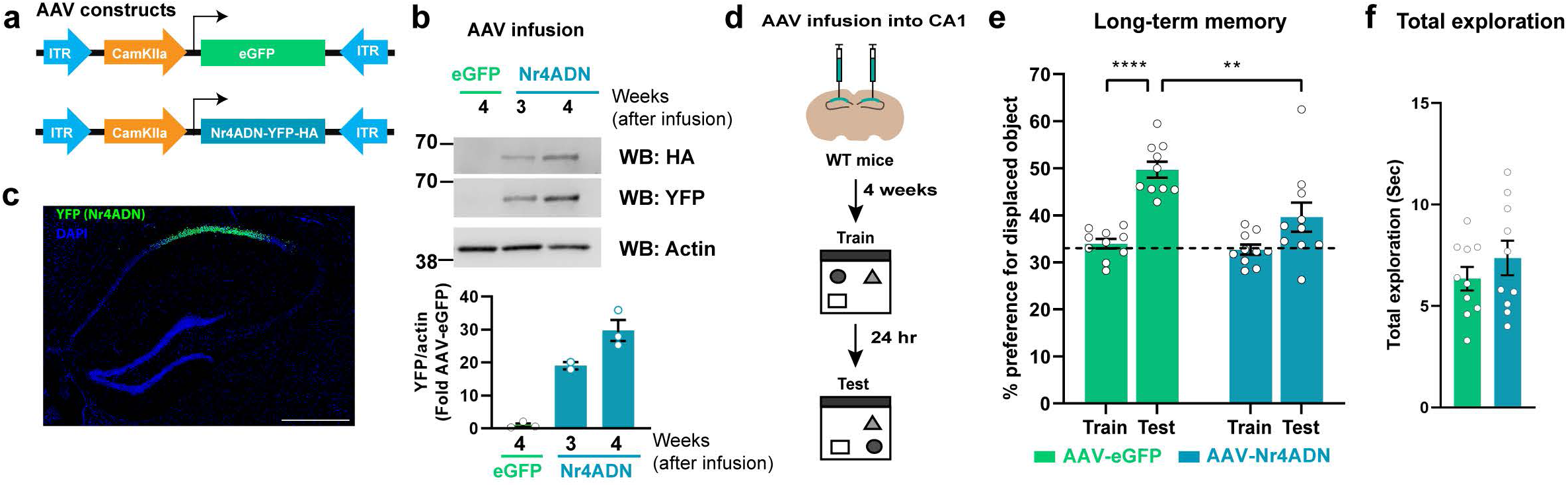
Functional validation of spatially reserved signatures of learning-induced gene expression. **a.** Design of the constructs packaged into Adeno-associated viruses (AAV) to ectopically express the dominant negative (DN) mutant of Nr4a and EGFP in the CA1 hippocampal sub-region. **b.** Western Blot analysis showing the time course of viral expression at 3-weeks and 4-weeks after viral infusion. One-way Anova: Šídák’s multiple comparisons test: eGFP vs Nr4ADN. n=2-3/group. **c.** Immunohistochemistry against YFP to detect the localization and spread of the AAV in the dorsal hippocampus. **d.** Experimental timeline of AAV-infusion into CA1 excitatory neurons followed by spatial learning paradigm. **e.** Long-term memory assessment by evaluating preference for the displaced object (DO) in a spatial object recognition (SOR) task. 2-way Anova: Significant sessions (Train-Test) x virus (Nr4ADN-eGFP) interaction: F (1, 18) = 4.537, p=0.0472, main effect of sessions: F (1, 18) = 29.93, p<0.0001 and main effect of virus: F (1, 18) = 10.26, p=0.0049. Šídák’s multiple comparisons test: eGFP: train vs test: p<0.0001, eGFP (test) vs Nr4ADN (Test): p=0.0014. n=10/group **f.** Total exploration time of all the objects during SOR for both the experimental groups.

Interestingly, protein kinase *Sgk1* was the only upregulated gene appearing in both the stratum radiatum and oriens but not in the CA1 pyramidal layer (**Fig 3f-g**). Distinct upregulation of *Sgk1* within stratum radiatum and oriens could be from interneurons or non-neuronal cells or displayed in this region due to the dendritic transport of mRNA from the CA1 pyramidal neurons. Similarly, *Tsc22d3* was found to be specifically induced in stratum radiatum, while *Rasgrp1* was exclusively induced in stratum oriens. Thus, using spatial transcriptomics, we can begin to understand how RNA is localized to subcellular compartments as a method of transcriptomic regulation, while this is unavailable from bulk and single nuclei transcriptomic datasets.

Although fewer genes were downregulated following learning compared to upregulated genes, *Kcna4, Usp2,* and *Shisa4* were downregulated in both CA1 and DG subregions (**Fig 3h**). *Kcna4* (Potassium Voltage-Gated Channel Subfamily A Member 4) expression was found to be increased in Abeta-induced cognitive impairment ^65^, while *Usp2* was found to be downregulated in hippocampus following sleep deprivation ^66^. As both sleep deprivation ^67^ and Abeta causes hippocampal memory deficits ^68^, altered expression of these genes indicate they have a possible role in learning and memory. Similarly, genes encoding two evolutionarily conserved RNA-binding proteins, *Rbm3* and *Cirbp,* were exclusively downregulated in DG (**Fig 3h**) and shown to be differentially expressed in the hippocampus following sleep deprivation ^66,69^. Among the genes downregulated exclusively in CA1 stratum oriens, *Mbp*, *Mobp* and *Plp1* are associated with structural constituents of myelin sheath, and *Opalin* is involved in oligodendrocyte differentiation. While adult oligodendrogenesis and myelination in the cortex are required for memory consolidation ^70^, the role of downregulation of these oligodendrocyte related genes in hippocampal subregion CA1 stratum oriens is not clear.

### Functional validation of spatial transcriptomic findings by subregion-specific manipulation of gene expression

The nuclear receptor 4a (Nr4a) subfamily of transcription factors are critical mediators of memory consolidation. They are robustly upregulated in the hippocampus within minutes after learning to regulate downstream gene expression ^57,71,72^. We have previously generated a dominant negative mouse model of Nr4a transcription factors which expresses a mutant form of Nr4a1 (Nr4ADN) lacking a key transcriptional activation domain ^58^ blocking downstream gene expression of all the Nr4a subfamily members ^73^. Our spatial transcriptomics data revealed upregulation of all the three members of the Nr4a subfamily (*Nr4a1*, *Nr4a2* and *Nr4a3*) in the CA1 pyramidal layer following learning (**Fig 3**). This signature was absent in other hippocampal subregions. Previous reports suggest that selectively knocking down the expression of either *Nr4a1* or *Nr4a2* in CA1 impairs spatial memory ^72^. Therefore, we sought to understand whether blocking the transcriptional activation function of all the three Nr4a family members exclusively in CA1 excitatory neurons would impair long-term memory consolidation. We used an adeno-associated viral construct of Nr4ADN (AAV-Nr4ADN; 2/2 stereotype to enable minimum diffusion across different subregions) under a CaMKIIα promoter to restrict expression to only excitatory neurons in CA1 (**Fig 4a, b and c**). To determine whether local expression of Nr4ADN in CA1 affects memory, AAV-Nr4ADN or control (AAV-eGFP) was infused into the dorsal CA1 of wild-type mice 4 weeks before SOR training (**Fig 4d**). Control mice showed a significant increase in preference for the displaced object during the 24 hr SOR test session relative to training, while AAV-Nr4ADN mice failed to show a preference for the displaced object (**Fig 4e**), demonstrating a long-term memory impairment in Nr4ADN expressing mice. Total exploration of the objects during the test session was unchanged and did not affect preference for the displaced object (**Fig 5f**). This finding functionally validates our spatial transcriptomics data; blocking Nr4a transcriptional function exclusively within CA1 excitatory neurons was sufficient to impair long-term memory.

## Discussion

In this study, we uncover a precise transcriptomic signature exhibited by different hippocampal subregions at a critical early timepoint during memory consolidation. While previous work has focused on studying gene expression changes in the whole hippocampus ^38,39,56,74^ and individual subregions ^40-42,44^, our study provides the first comprehensive analysis of the simultaneous transcriptomic changes spatially distributed across the hippocampal subregions in response to learning. Moreover, we functionally validated spatial transcriptomic analyses demonstrating that blocking the activity of Nr4a subfamily of transcription factors selectively within CA1 leads to long-term memory deficits.

Within the dorsal hippocampus, the CA1 pyramidal layer, stratum radiatum, and stratum oriens are critical for encoding spatial memory ^75^ . While these principal layers plays a role in generating spatial maps of the environment ^7,46^, the granule cells within the DG are thought to provide stable representations of a specific environment ^76–78^. In this study, we identified differential expression patterns for some of the most extensively studied IEGs related to transcriptional regulation in the CA principal layers (CA1 and CA2/3) after spatial exploration. *Nr4a1* and *Egr3* were predominantly induced in CA1 subregions, whereas *Arc*, *Nr4a2*, *Per1,* and *Dusp5* were upregulated in CA1 and CA2/3 regions. IEGs *Egr1* and *Homer1* were found to be upregulated in all sub-regions studied, while *Gadd45b* and *Per2* were induced exclusively in DG. Differential gene induction has been correlated with activation of engram ensembles ^4–7^ and place codes underlying spatial maps ^7,46^. We also noted a greater number of differentially expressed genes in DG compared to CA1 following spatial exploration, consistent with single nuclei data from activated and non-activated neurons from DG and CA1 ^48^, although we found that the CA1 subregion exhibited a greater number of IEGs associated with activated engram ensembles ^5–7^. Additionally, our study highlights transcriptomic signatures within the two relatively understudied hippocampal compartments, stratum radiatum and stratum oriens, which have been challenging to delineate using conventional single-cell sequencing strategies. Overall, our study elucidates the transcriptomic diversity that prevails between hippocampal subregions during an early window of spatial memory consolidation.

The nuclear receptor 4a (Nr4a) subfamily, *Nr4a1, Nr4a2,* and *Nr4a3*, serve as major regulators of gene expression in the hippocampus during memory consolidation ^58,59,72,73,79,80^. *Nr4a1* has been implicated in regulating object location memory, while both *Nr4a1* and *Nr4a2* are necessary for object location and object recognition memory in the dorsal hippocampus ^72^. Impairments in Nr4a function ^58,81^ leads to long-term memory deficits ^57,58^ and impairments in transcription-dependent long-term potentiation (LTP) in CA1 ^82^. On the contrary, overexpression or pharmacological activation of Nr4a family members ameliorates memory deficits in mouse models of Alzheimer’s disease and related dementias (ADRD) and age-associated memory decline ^58,59,71,83^. Our identification of increased expression of *Nr4a* subfamily members after learning in CA1 confirms findings from previous studies using hippocampus-dependent learning tasks ^71,72,84,85^. Further, we validated our findings by demonstrating the functional relevance of CA1-specific *Nr4a* expression in long-term spatial memory. Thus, integrating the spatial component of learning-induced transcriptomic heterogeneity in the hippocampal cell layers strongly supports the concept of subregion-specific dissociation in the molecular mechanisms underlying memory consolidation.

The basal dendrites of CA1 pyramidal neurons make up stratum oriens while stratum radiatum consists of apical dendrites. Both stratum radiatum and oriens receive inputs from CA3 Schaffer collaterals ^86^. We found upregulation in *Nr4a1*, *Homer1*, *Egr1*, *Egr3*, *Egr4, Dnajb5, and Hspa5* in the CA1 pyramidal layer, CA1 stratum radiatum and oriens. Interestingly, *Sgk1* was restricted only to the stratum oriens and stratum radiatum-suggesting *Sgk1* could be enriched in the dendritic region to enable local translation of this regulatory kinase in response to synaptic activity ^87^. However, interneuron and non-neuronal cells within stratum radiatum and oriens layers could also exhibit learning-induced upregulation of *Sgk1*. Importantly, Sgk1 plays a functional role in memory consolidation. Expression of a dominant negative Sgk1 within CA1 impaired spatial memory ^32^, whereas constitutively active Sgk1 enhanced spatial memory ^31^. Furthermore, in an APP/PS1 based ADRD model, *Sgk1* was downregulated in the hippocampus, whereas overexpression of Sgk1 could ameliorate spatial memory deficits ^34^. Sgk1 regulates the expression of *zif268/Egr1* ^88^, an IEG that we found upregulated in all subregions of the hippocampus following learning. Studying the spatial patterns of learning-responsive genes like *Sgk1* helps us define the role of specific hippocampal subregions in memory consolidation.

Our study has identified two upregulated pathways in DG that are involved in protein kinase inhibitor activity and protein processing in the endoplasmic reticulum (ER). We have recently shown that learning induces the expression of molecular chaperones localized at the ER, and this protein folding machinery is critical in synaptic plasticity and long-term memory consolidation ^58^. Here, our spatial transcriptomics data shows upregulation of genes encoding chaperones in distinct subregions; *Hspa5* and *Dnajb5* across all the hippocampal subregions, *Xbp1*, *Sdf2l1* and *Dnajb1* in areas CA1 and DG, and *Pdia6* and *Creld2* exclusively in DG. This suggests that DG could have a prominent role in ER protein processing during an early timepoint after spatial learning; although global upregulation of ER chaperones across all the subregions supports our previous findings that ER chaperones are indeed critical mediators of long-term memory storage ^58^. This work also suggests that there may be distinct protein processing complexes in different hippocampal subregions during memory consolidation that may be involved in the folding and surface presentation of distinct proteins.

Our work demonstrates that the subregions of the dorsal hippocampus respond to learning by exhibiting distinct transcriptomic signatures. These subregions differ by their circuitry, cell types, and electrophysiological features. However, a criticism of this spatial transcriptomic approach is that it lacks cell-type specific information, yet we see changes in some non-neuronal genes after learning. Therefore, future studies will need to address heterogeneity between cell types and how each of them responds to learning. Thus, combining spatial transcriptomics with single-cell transcriptomics and high throughput *in situ* approaches such as MERFISH ^89^ will provide further insights into cell-type specific changes in gene expression across different hippocampal sub-regions during memory consolidation. Although this study focused on spatial transcriptomic signatures at an early critical time-window of spatial learning, the differential gene expression patterns we identified may well lead to diverse profiles of target gene activation across brain regions at later timepoints. Our attempt to elucidate the spatial transcriptomic signature of memory provides the groundwork for future studies to understand the precise gene expression patterns underlying memory consolidation, and whether these signatures are affected in neurological disorders associated with memory impairments.

## Acknowledgments

Spatial gene expression using the *10X Genomics Visium* platform was performed at the Iowa NeuroBank Core in the Iowa Neuroscience Institute, and the Genomics Division in the Iowa Institute of Human Genetics which is supported, in part, by the University of Iowa Carver College of Medicine. We thank the Neural Circuits and Behavior Core at the Iowa Neuroscience Institute for use of their facilities. We thank Emily N. Walsh for technical assistance, Dr. Mahesh S. Shetty and Dr. Lisa Lyons for advice on the manuscript. We thank Xiaowen Wang (Partek Inc.) for technical support which was crucial for Visium spatial gene expression data analysis.

## Author contributions

SC and TA conceived the idea. SC, LL and UM wrote the manuscript with inputs from all the authors. SC performed viral infusion, behavior, biochemical and molecular biology experiments, analyzed and interpreted the data. YV performed all the bioinformatic analysis, generated plots and analyzed the data with inputs from EB and JJM. LCL processed the tissue for Visium. UM performed imaging, UM and LL performed biochemical experiments.

## Materials and methods

### Data reporting

No statistical methods were used to predetermine sample size.

### Mouse lines

Adult male C57BL/6J mice were purchased from Jackson Laboratories were 2-3 months age during behavioral or biochemical experiments. All mice had free access to food and water, and lights were maintained on 12h light/dark cycle. All behavioral testing was performed during the light cycle between Zeitgeber time (ZT) 0-2. For all behavioral and biochemical experiments, mice were randomly assigned to groups, were house individually, and were handled for 2 min per day for 5 days. All experiments were conducted according to US National Institutes of Health guidelines for animal care and use and were approved by the Institutional Animal Care and Use Committee of the University of Iowa, Iowa.

### Adeno-associated virus (AAV) constructs and stereotactic surgeries

AAV_2.2_-CaMKIIα-Nr4ADN and AAV_2.2_-CaMKIIα-EGFP were purchased from VectorBuilder (VectorBuilder Inc). Stereotactic surgeries were performed as previously described ^58^. Briefly, mice were anaesthetized using isoflurane and 1 μl of respective AAVs were injected into the dorsal hippocampus (coordinates: anteroposterior, −1.9 mm, mediolateral, ±1.5 mm, and 1.5 mm below bregma). Following viral infusion, drill holes were closed with bone wax (Lukens) and the incisions were sutured.

### Spatial object recognition (SOR) task

SOR was performed as previously described ^58^. Animals were handled for 5 consecutive days before training. On the day of training, animals were briefly habituated in an open field, followed by three 6-minute sessions inside an arena containing three different objects. 24 hr later, the animals were returned to the arena with one of the objects displaced to a novel spatial coordinate. Exploration time around all the objects were then manually scored.

### Visium sample preparation

After rapidly euthanized by cervical dislocation, the brains from 8 mice were rapidly extracted and flash- frozen with −70°C isopentane for 5 minutes. Frozen brains were stored at −80C until sectioning. Mouse frozen brains were embedded in optimal cutting temperature medium (OCT) and cryosectioned at −20 °C with the Leica CM3050 S Cryostat. 10-microns of coronal sections from the brain region with dorsal hippocampus were placed on chilled Visium Tissue Optimization Slides (10X Genomics) and Visium Spatial Gene Expression Slides (10X Genomics). Visium slides with the sections were fixed, stained, and imaged with Hematoxylin and Eosin using a 20X objective on an Olympus BX61 Upright Microscope. Tissue was then permeabilized for 18 min, which was established an optimal permeabilization time based on tissue optimization time-course experiments. The poly-A mRNAs from the slices were released and captured by the poly(dT) primers and precoated on the slide, including a spatial barcode and a Unique Molecular Identifiers (UMIs). After reverse transcription and second strand synthesis, the amplificated cDNA samples from the Visium slides were transferred, purified, and quantified for library preparation. The fragmented cDNA samples were used to construct sequencing for Visium spatial transcriptome on a NovaSeq 6000 (Illumina) at a sequencing depth of 150 million total read pairs per mouse Visium sample.

### Visium library preparation and sequencing

Sequencing libraries were prepared by the Iowa Institute of Human Genetics (IIHG) Genomics Division, according to the Visium Spatial Gene Expression User Guide. Each pooled library was sequenced on an Illumina NovaSeq 6000 using SBS chemistry v1.5 for 100 cycles, at a sequencing depth of 200 million total read pairs. Data processing of Visium data, raw FASTQ files and images were output with Space Ranger software (Version 1.3.1) and analyzed downstream by Partek Flow (Partek Inc.) with their single-cell analysis pipeline, mm10 reference genome was used for gene alignment.

### Visium data analysis

The read counts were normalized by the counts per million (CPM) method and transformed to log2(CPM + 1). A general linear model was applied to correct for batch effect between the two sets of experiment. Hippocampal subregions were selected based on biological knowledge using anatomical structures apparent on the H&E staining images. The pyramidal layers of CA1, CA2+CA3 and granular and molecular layer of DG were selected for their role in neuronal excitability, synaptic plasticity and memory. Additionally, CA1 stratum radiatum and oriens were also selected due to their roles in neuronal circuitry. Differential gene expression analysis was performed using the non-parametric Kruskal-Wallis rank sum test because this type of tests have been the most widely used approach in the field of single-cell transcriptomics (Squair et al. 2021). Because each cell is assumed to be a biological replicate in scRNA-seq, the same assumption is made here for each visium spot which generates a big sample size that is handled correctly by Kruskal-Wallis test. Gene-specific analyses were filtered with false discovery rate (FDR) < 0.05 and fold change > |1.4|.

### Bulk RNA extraction, cDNA preparation and gene expression analysis

Dorsal hippocampi were dissected and immediately stored at −80°C in RNAlater solution (Ambion). For RNA total extraction, hippocampi were homogenized in Qiazol (Qiagen) using stainless steel beads (Qiagen). Chloroform was then added, and the homogenate was centrifuged at 12,000 x g at RT for 15 min. Aqueous phase containing RNA was precipitated using ethanol and then cleaned using the RNeasy kit (Qiagen). RNA was eluted in nuclease-free water, treated with DNase (Qiagen) at RT for 25 min and precipitated in ethanol, sodium acetate (pH 5.2) and glycogen overnight at −20°C. Precipitated RNA samples were centrifuged at top speed at RT for 20 min, washed with 70% ethanol and centrifuged at top speed for 5 min, dried and resuspended in nuclease free water. RNA concentrations were estimated using a Nanodrop (Thermo Fisher Scientific). cDNAs were prepared from 1 μg RNA using the SuperScript™ IV First-Strand Synthesis System (Ambion). Real-time RT-PCR reactions were performed in a 384-well optical reaction plate with optical adhesive covers (Life Technologies). Each reaction was composed of 2.25μl cDNA (2 ng/ul), 2.5μl Fast SYBR™ Green Master Mix (Thermo Fisher Scientific), and 0.25μl of primer mix (IDT). Three technical replicates per reaction was performed on the QuantStudio 7 Flex Real-Time PCR system (Applied Biosystems, Life Technologies). Data was normalized to housekeeping genes (Tubulin, Pgk1 and Hprt) and 2^(-ΔΔCt)^ method was used for gene expression analysis.

### Library preparation and sequencing from bulk RNA

RNA libraries were prepared at the Iowa Institute of Human Genetics (IIHG), Genomics Division, using the Illumina TruSeq Stranded Total RNA with Ribo-Zero gold sample preparation kit (Illumina, Inc., San Diego, CA). Library concentrations were measured using KAPA Illumina Library Quantification Kit (KAPA Biosystems, Wilmington, MA). Polled libraries were sequenced on Illumina NovaSeq6000 sequencer with 150-bp Paired-End chemistry (Illumina) at the IIHG core.

### Bulk RNA-seq analysis

Sequencing data was processed with the bcbio-nextgen pipeline (https://github.com/bcbio/bcbio-nextgen). The pipeline uses STAR ^90^ to align reads to the genome and quantifies expression at the gene level with featureCounts ^91^. All further analyses were performed using R. For gene level count data, the R package EDASeq was used to account for sequencing depth (upper quartile normalization) ^92^. Latent sources of variation in expression levels were assessed and accounted for using RUVSeq (RUVs) ^93^. Appropriate choice of the RUVSeq parameter k was determined through inspection of RLE plots and PCA plots. Differential expression analysis was conducted using edgeR ^94^.

## Molecular function enrichment analysis

The identified DEGs were analyzed for molecular function enrichment analysis by using the ClueGO and CluePedia plug-ins of the Cytoscape 3.9.0 software in “Functional analysis” mode against the Gene Ontology Molecular Function (4691 terms) database. The GO Tern Fusion was used allowing for the fusion of GO parent-child terms based on similar associated genes. The GO Term Connectivity had a kappa score of 0.4. The enrichment was performed using a two-sided hypergeometric test. The p-values were corrected with a Bonferroni step down approach. Only significant molecular function with corrected p-values < 0.05 were displayed. UpSet plots were generated using an online software ExpressAnalyst. Data was plotted using the distinct mode.

### Western blot analysis

Protein extracts were transferred to polyvinylidene difluoride membranes as previously described ^74^. Membranes were blocked with Odyssey® Blocking Buffer in TBS (LI-COR) and incubated overnight at 4°C with the following primary antibodies: pan-HA (1:1000, Cell signaling), YFP (1:1000, Abcam), and Actin (1:10,000, ThermoFisher Scientific). Membranes were washed and incubated with appropriate IRDye IgG secondary antibodies, including anti-rabbit IRDye 800LT (1:5,000, LI-COR) and anti-mouse IRDye 680CW (LI-COR). Images were acquired using the Odyssey Infrared Imaging System (LI-COR). Quantification of western blot bands was performed using Image Studio Lite ver5.2 (LI-COR).

### Immunohistochemistry and confocal imaging

Animals were perfused with 4% PFA, and 20 μm coronal brain sections were made in a cryostat. Free-floating sections were washed with PBS and mounted on on Superfrost™ Plus microscope slides (Fisherbrand). The sections were air-dried, followed by coverslip mounting with Vectashield® Antifade Mounting Medium with DAPI (Vector Laboratories). Slides were then imaged using the Olympus FV3000 confocal microscope with a 10X NA = 0.4 objective at 800 × 800-pixel resolution.

### Statistics

Behavioral and biochemical data were analyzed using unpaired two-tailed t-tests and either one-way or two-way ANOVAs (in some cases with repeated measures as the within subject variable). Sidak’s tests were used for post-hoc analyses where needed. Differences were considered statistically significant when p<0.05. As indicated for each figure panel, all data are plotted in either bar graphs, in which symbols represent each data point, or in dot plots, where each symbol represents an individual data point. Graphs were plotted as mean ± SEM.

## Graphical abstract

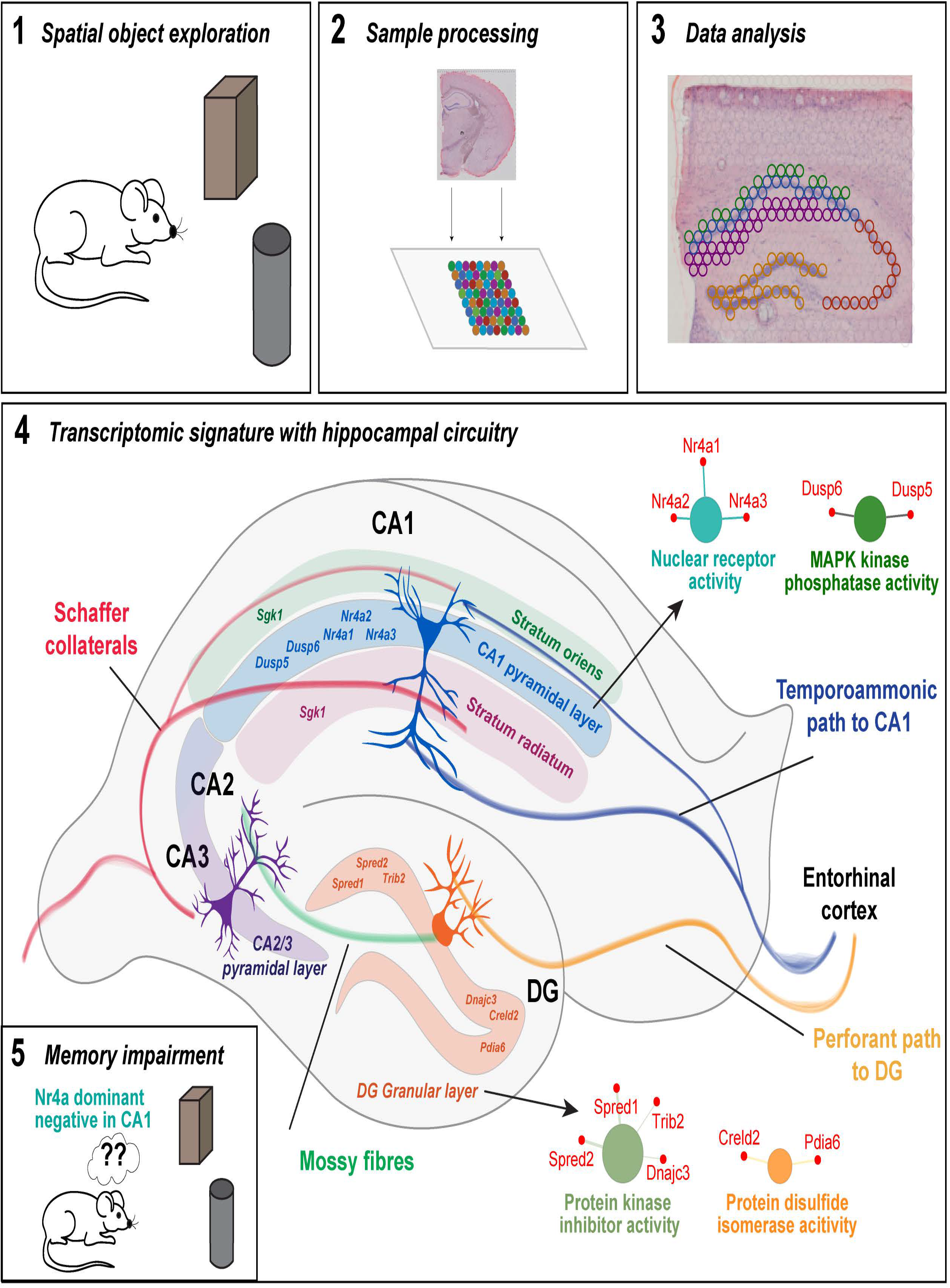

## References

1 Yap, E. L. & Greenberg, M. E. Activity-Regulated Transcription: Bridging the Gap between Neural Activity and Behavior. Neuron 100, 330–348 (2018). https://doi.org:10.1016/j.neuron.2018.10.013

2 Fernandez-Albert, J. et al. Immediate and deferred epigenomic signatures of in vivo neuronal activation in mouse hippocampus. Nat Neurosci 22, 1718–1730 (2019). https://doi.org:10.1038/s41593-019-0476-2

3 Tyssowski, K. M. et al. Different Neuronal Activity Patterns Induce Different Gene Expression Programs. Neuron 98, 530–546 e511 (2018). https://doi.org:10.1016/j.neuron.2018.04.001

4 Roy, D. S. et al. Brain-wide mapping reveals that engrams for a single memory are distributed across multiple brain regions. Nat Commun 13, 1799 (2022). https://doi.org:10.1038/s41467-022-29384-4

5 Josselyn, S. A. & Tonegawa, S. Memory engrams: Recalling the past and imagining the future. Science 367 (2020). https://doi.org:10.1126/science.aaw4325

6 Reijmers, L. G., Perkins, B. L., Matsuo, N. & Mayford, M. Localization of a stable neural correlate of associative memory. Science 317, 1230–1233 (2007). https://doi.org:10.1126/science.1143839

7 Pettit, N. L., Yap, E. L., Greenberg, M. E. & Harvey, C. D. Fos ensembles encode and shape stable spatial maps in the hippocampus. Nature 609, 327–334 (2022). https://doi.org:10.1038/s41586-022-05113-1

8 Redondo, R. L. et al. Bidirectional switch of the valence associated with a hippocampal contextual memory engram. Nature 513, 426–430 (2014). https://doi.org:10.1038/nature13725

9 Cowansage, K. K. et al. Direct reactivation of a coherent neocortical memory of context. Neuron 84, 432–441 (2014). https://doi.org:10.1016/j.neuron.2014.09.022

10 Kitamura, T. et al. Engrams and circuits crucial for systems consolidation of a memory. Science 356, 73–78 (2017). https://doi.org:10.1126/science.aam6808

11 Guzowski, J. F., McNaughton, B. L., Barnes, C. A. & Worley, P. F. Environment-specific expression of the immediate-early gene Arc in hippocampal neuronal ensembles. Nat Neurosci 2, 1120–1124 (1999). https://doi.org:10.1038/16046

12 Tonegawa, S., Liu, X., Ramirez, S. & Redondo, R. Memory Engram Cells Have Come of Age. Neuron 87, 918–931 (2015). https://doi.org:10.1016/j.neuron.2015.08.002

13 Liu, X. et al. Optogenetic stimulation of a hippocampal engram activates fear memory recall. Nature 484, 381–385 (2012). https://doi.org:10.1038/nature11028

14 Han, J. H. et al. Selective erasure of a fear memory. Science 323, 1492–1496 (2009). https://doi.org:10.1126/science.1164139

15 Broadbent, N. J., Squire, L. R. & Clark, R. E. Spatial memory, recognition memory, and the hippocampus. Proc Natl Acad Sci U S A 101, 14515–14520 (2004). https://doi.org:10.1073/pnas.0406344101

16 Moser, M. B., Moser, E. I., Forrest, E., Andersen, P. & Morris, R. G. Spatial learning with a minislab in the dorsal hippocampus. Proc Natl Acad Sci U S A 92, 9697–9701 (1995). https://doi.org:10.1073/pnas.92.21.9697

17 Moser, M. B. & Moser, E. I. Functional differentiation in the hippocampus. Hippocampus 8, 608–619 (1998). https://doi.org:10.1002/(SICI)1098-1063(1998)8:6<608::AID-HIPO3>3.0.CO;2-7

18 Hainmueller, T. & Bartos, M. Dentate gyrus circuits for encoding, retrieval and discrimination of episodic memories. Nat Rev Neurosci 21, 153–168 (2020). https://doi.org:10.1038/s41583-019-0260-z

19 Witter, M. P., Griffioen, A. W., Jorritsma-Byham, B. & Krijnen, J. L. Entorhinal projections to the hippocampal CA1 region in the rat: an underestimated pathway. Neurosci Lett 85, 193–198 (1988). https://doi.org:10.1016/0304-3940(88)90350-3

20 Deller, T., Adelmann, G., Nitsch, R. & Frotscher, M. The alvear pathway of the rat hippocampus. Cell Tissue Res 286, 293–303 (1996). https://doi.org:10.1007/s004410050699

21 Vago, D. R. & Kesner, R. P. Disruption of the direct perforant path input to the CA1 subregion of the dorsal hippocampus interferes with spatial working memory and novelty detection. Behav Brain Res 189, 273–283 (2008). https://doi.org:10.1016/j.bbr.2008.01.002

22 Remondes, M. & Schuman, E. M. Role for a cortical input to hippocampal area CA1 in the consolidation of a long-term memory. Nature 431, 699–703 (2004). https://doi.org:10.1038/nature02965

23 Place, R. et al. NMDA signaling in CA1 mediates selectively the spatial component of episodic memory. Learn Mem 19, 164–169 (2012). https://doi.org:10.1101/lm.025254.111

24 Huerta, P. T., Sun, L. D., Wilson, M. A. & Tonegawa, S. Formation of temporal memory requires NMDA receptors within CA1 pyramidal neurons. Neuron 25, 473–480 (2000). https://doi.org:10.1016/s0896-6273(00)80909-5

25 Steward, O. Topographic organization of the projections from the entorhinal area to the hippocampal formation of the rat. J Comp Neurol 167, 285–314 (1976). https://doi.org:10.1002/cne.901670303

26 Arrigoni, E. & Greene, R. W. Schaffer collateral and perforant path inputs activate different subtypes of NMDA receptors on the same CA1 pyramidal cell. Br J Pharmacol 142, 317–322 (2004). https://doi.org:10.1038/sj.bjp.0705744

27 Anand, K. S. & Dhikav, V. Hippocampus in health and disease: An overview. Ann Indian Acad Neurol 15, 239–246 (2012). https://doi.org:10.4103/0972-2327.104323

28 Amaral David, L. P. in The Hippocampus Book (ed Richard Morris Per Andersen, David Amaral, Tim Bliss, John O’Keefe) Ch. 3, (Oxford University Press, 2006).

29 Drew, L. J., Fusi, S. & Hen, R. Adult neurogenesis in the mammalian hippocampus: why the dentate gyrus? Learn Mem 20, 710–729 (2013). https://doi.org:10.1101/lm.026542.112

30 McHugh, T. J. et al. Dentate gyrus NMDA receptors mediate rapid pattern separation in the hippocampal network. Science 317, 94–99 (2007). https://doi.org:10.1126/science.1140263

31 Nakazawa, K. et al. Requirement for hippocampal CA3 NMDA receptors in associative memory recall. Science 297, 211–218 (2002). https://doi.org:10.1126/science.1071795

32 Nakashiba, T. et al. Young dentate granule cells mediate pattern separation, whereas old granule cells facilitate pattern completion. Cell 149, 188–201 (2012). https://doi.org:10.1016/j.cell.2012.01.046

33 Stevenson, R. F., Reagh, Z. M., Chun, A. P., Murray, E. A. & Yassa, M. A. Pattern Separation and Source Memory Engage Distinct Hippocampal and Neocortical Regions during Retrieval. J Neurosci 40, 843–851 (2020). https://doi.org:10.1523/JNEUROSCI.0564-19.2019

34 Komorowski, R. W., Manns, J. R. & Eichenbaum, H. Robust conjunctive item-place coding by hippocampal neurons parallels learning what happens where. J Neurosci 29, 9918–9929 (2009). https://doi.org:10.1523/JNEUROSCI.1378-09.2009

35 Dimsdale-Zucker, H. R., Ritchey, M., Ekstrom, A. D., Yonelinas, A. P. & Ranganath, C. CA1 and CA3 differentially support spontaneous retrieval of episodic contexts within human hippocampal subfields. Nat Commun 9, 294 (2018). https://doi.org:10.1038/s41467-017-02752-1

36 Favila, S. E., Chanales, A. J. & Kuhl, B. A. Experience-dependent hippocampal pattern differentiation prevents interference during subsequent learning. Nat Commun 7, 11066 (2016). https://doi.org:10.1038/ncomms11066

37 Schlichting, M. L., Mumford, J. A. & Preston, A. R. Learning-related representational changes reveal dissociable integration and separation signatures in the hippocampus and prefrontal cortex. Nat Commun 6, 8151 (2015). https://doi.org:10.1038/ncomms9151

38 Poplawski, S. G. et al. Contextual fear conditioning induces differential alternative splicing. Neurobiol Learn Mem 134 Pt B, 221–235 (2016). https://doi.org:10.1016/j.nlm.2016.07.018

39 Peixoto, L. L. et al. Memory acquisition and retrieval impact different epigenetic processes that regulate gene expression. BMC Genomics 16 **Suppl** 5, S5 (2015). https://doi.org:10.1186/1471-2164-16-S5-S5

40 Benito, E. et al. HDAC inhibitor-dependent transcriptome and memory reinstatement in cognitive decline models. J Clin Invest 125, 3572–3584 (2015). https://doi.org:10.1172/JCI79942

41 Halder, R. et al. DNA methylation changes in plasticity genes accompany the formation and maintenance of memory. Nat Neurosci 19, 102–110 (2016). https://doi.org:10.1038/nn.4194

42 Gregoire, C. A. et al. RNA-Sequencing Reveals Unique Transcriptional Signatures of Running and Running-Independent Environmental Enrichment in the Adult Mouse Dentate Gyrus. Front Mol Neurosci 11, 126 (2018). https://doi.org:10.3389/fnmol.2018.00126

43 Chen, P. B. et al. Mapping Gene Expression in Excitatory Neurons during Hippocampal Late-Phase Long-Term Potentiation. Front Mol Neurosci 10, 39 (2017). https://doi.org:10.3389/fnmol.2017.00039

44 Lacar, B. et al. Nuclear RNA-seq of single neurons reveals molecular signatures of activation. Nat Commun 7, 11022 (2016). https://doi.org:10.1038/ncomms11022

45 Marco, A. et al. Mapping the epigenomic and transcriptomic interplay during memory formation and recall in the hippocampal engram ensemble. Nat Neurosci 23, 1606–1617 (2020). https://doi.org:10.1038/s41593-020-00717-0

46 O’Keefe, J. Place units in the hippocampus of the freely moving rat. Exp Neurol 51, 78–109 (1976). https://doi.org:10.1016/0014-4886(76)90055-8

47 Yap, E. L. et al. Bidirectional perisomatic inhibitory plasticity of a Fos neuronal network. Nature 590, 115–121 (2021). https://doi.org:10.1038/s41586-020-3031-0

48 Jaeger, B. N. et al. A novel environment-evoked transcriptional signature predicts reactivity in single dentate granule neurons. Nat Commun 9, 3084 (2018). https://doi.org:10.1038/s41467-018-05418-8

49 Hrvatin, S. et al. Single-cell analysis of experience-dependent transcriptomic states in the mouse visual cortex. Nat Neurosci 21, 120–129 (2018). https://doi.org:10.1038/s41593-017-0029-5

50 Wu, Y. E., Pan, L., Zuo, Y., Li, X. & Hong, W. Detecting Activated Cell Populations Using Single-Cell RNA-Seq. Neuron 96, 313–329 e316 (2017). https://doi.org:10.1016/j.neuron.2017.09.026

51 Maynard, K. R. et al. Transcriptome-scale spatial gene expression in the human dorsolateral prefrontal cortex. Nat Neurosci 24, 425–436 (2021). https://doi.org:10.1038/s41593-020-00787-0

52 Farris, S. et al. Hippocampal Subregions Express Distinct Dendritic Transcriptomes that Reveal Differences in Mitochondrial Function in CA2. Cell Rep 29, 522–539 e526 (2019). https://doi.org:10.1016/j.celrep.2019.08.093

53 Chen, W. T. et al. Spatial Transcriptomics and In Situ Sequencing to Study Alzheimer’s Disease. Cell 182, 976–991 e919 (2020). https://doi.org:10.1016/j.cell.2020.06.038

54 Bahl E, C. S., Elsadany M, Vanrobaeys Y, Lin L-C, Giese KP, Abel T, Michaelson JJ. NEUROeSTIMator: Using Deep Learning to Quantify Neuronal Activation from Single-Cell and Spatial Transcriptomic Data. bioRxiv (2022). https://doi.org:https://doi.org/10.1101/2022.04.08.487573

55 Peixoto, L. L. et al. Memory acquisition and retrieval impact different epigenetic processes that regulate gene expression. BMC Genomics 16 **Suppl** 5, S5 (2015). https://doi.org:10.1186/1471-2164-16-S5-S5

56 Chatterjee, S. et al. The CBP KIX domain regulates long-term memory and circadian activity. BMC Biol 18, 155 (2020). https://doi.org:10.1186/s12915-020-00886-1

57 Hawk, J. D. et al. NR4A nuclear receptors support memory enhancement by histone deacetylase inhibitors. J Clin Invest 122, 3593–3602 (2012). https://doi.org:10.1172/JCI64145

58 Chatterjee, S. et al. Endoplasmic reticulum chaperone genes encode effectors of long-term memory. Sci Adv 8, eabm6063 (2022). https://doi.org:10.1126/sciadv.abm6063

59 Chatterjee, S. et al. Pharmacological activation of Nr4a rescues age-associated memory decline. Neurobiol Aging 85, 140–144 (2020). https://doi.org:10.1016/j.neurobiolaging.2019.10.001

60 Kim, H. J. et al. Histone demethylase PHF2 activates CREB and promotes memory consolidation. EMBO Rep 20, e45907 (2019). https://doi.org:10.15252/embr.201845907

61 Saez, M. A. et al. Mutations in JMJD1C are involved in Rett syndrome and intellectual disability. Genet Med 18, 378–385 (2016). https://doi.org:10.1038/gim.2015.100

62 Phoenix, T. N. & Temple, S. Spred1, a negative regulator of Ras-MAPK-ERK, is enriched in CNS germinal zones, dampens NSC proliferation, and maintains ventricular zone structure. Genes Dev 24, 45–56 (2010). https://doi.org:10.1101/gad.1839510

63 Abel, T., Martin, K. C., Bartsch, D. & Kandel, E. R. Memory suppressor genes: inhibitory constraints on the storage of long-term memory. Science 279, 338–341 (1998). https://doi.org:10.1126/science.279.5349.338

64 Kwapis, J. L. et al. Epigenetic regulation of the circadian gene Per1 contributes to age-related changes in hippocampal memory. Nat Commun 9, 3323 (2018). https://doi.org:10.1038/s41467-018-05868-0

65 Pan, Y., Xu, X., Tong, X. & Wang, X. Messenger RNA and protein expression analysis of voltage-gated potassium channels in the brain of Abeta(25-35)-treated rats. J Neurosci Res 77, 94–99 (2004). https://doi.org:10.1002/jnr.20134

66 Vecsey, C. G. et al. Genomic analysis of sleep deprivation reveals translational regulation in the hippocampus. Physiol Genomics 44, 981–991 (2012). https://doi.org:10.1152/physiolgenomics.00084.2012

67 Havekes, R. et al. Sleep deprivation causes memory deficits by negatively impacting neuronal connectivity in hippocampal area CA1. Elife 5 (2016). https://doi.org:10.7554/eLife.13424

68 Serneels, L. et al. gamma-Secretase heterogeneity in the Aph1 subunit: relevance for Alzheimer’s disease. Science 324, 639–642 (2009). https://doi.org:10.1126/science.1171176

69 Gaine, M. E. et al. Altered hippocampal transcriptome dynamics following sleep deprivation. Mol Brain 14, 125 (2021). https://doi.org:10.1186/s13041-021-00835-1

70 Steadman, P. E. et al. Disruption of Oligodendrogenesis Impairs Memory Consolidation in Adult Mice. Neuron 105, 150–164 e156 (2020). https://doi.org:10.1016/j.neuron.2019.10.013

71 Kwapis, J. L. et al. HDAC3-Mediated Repression of the Nr4a Family Contributes to Age-Related Impairments in Long-Term Memory. J Neurosci 39, 4999–5009 (2019). https://doi.org:10.1523/JNEUROSCI.2799-18.2019

72 McNulty, S. E. et al. Differential roles for Nr4a1 and Nr4a2 in object location vs. object recognition long-term memory. Learn Mem 19, 588–592 (2012). https://doi.org:10.1101/lm.026385.112

73 Hawk, J. D. & Abel, T. The role of NR4A transcription factors in memory formation. Brain Res Bull 85, 21–29 (2011). https://doi.org:10.1016/j.brainresbull.2011.02.001

74 Chatterjee, S. et al. Reinstating plasticity and memory in a tauopathy mouse model with an acetyltransferase activator. EMBO Mol Med 10 (2018). https://doi.org:10.15252/emmm.201708587

75 Grienberger, C., Milstein, A. D., Bittner, K. C., Romani, S. & Magee, J. C. Inhibitory suppression of heterogeneously tuned excitation enhances spatial coding in CA1 place cells. Nat Neurosci 20, 417–426 (2017). https://doi.org:10.1038/nn.4486

76 GoodSmith, D. et al. Spatial Representations of Granule Cells and Mossy Cells of the Dentate Gyrus. Neuron 93, 677–690 e675 (2017). https://doi.org:10.1016/j.neuron.2016.12.026

77 Senzai, Y. & Buzsaki, G. Physiological Properties and Behavioral Correlates of Hippocampal Granule Cells and Mossy Cells. Neuron 93, 691–704 e695 (2017). https://doi.org:10.1016/j.neuron.2016.12.011

78 Hainmueller, T. & Bartos, M. Parallel emergence of stable and dynamic memory engrams in the hippocampus. Nature 558, 292–296 (2018). https://doi.org:10.1038/s41586-018-0191-2

79 Safe, S. et al. Nuclear receptor 4A (NR4A) family - orphans no more. J Steroid Biochem Mol Biol 157, 48–60 (2016). https://doi.org:10.1016/j.jsbmb.2015.04.016

80 Bridi, M. S., Hawk, J. D., Chatterjee, S., Safe, S. & Abel, T. Pharmacological Activators of the NR4A Nuclear Receptors Enhance LTP in a CREB/CBP-Dependent Manner. Neuropsychopharmacology 42, 1243–1253 (2017). https://doi.org:10.1038/npp.2016.253

81 McQuown, S. C. et al. HDAC3 is a critical negative regulator of long-term memory formation. J Neurosci 31, 764–774 (2011). https://doi.org:10.1523/JNEUROSCI.5052-10.2011

82 Bridi, M. S. & Abel, T. The NR4A orphan nuclear receptors mediate transcription-dependent hippocampal synaptic plasticity. Neurobiol Learn Mem 105, 151–158 (2013). https://doi.org:10.1016/j.nlm.2013.06.020

83 Moon, M. et al. Nurr1 (NR4A2) regulates Alzheimer’s disease-related pathogenesis and cognitive function in the 5XFAD mouse model. Aging Cell 18, e12866 (2019). https://doi.org:10.1111/acel.12866

84 Pena de Ortiz, S., Maldonado-Vlaar, C. S. & Carrasquillo, Y. Hippocampal expression of the orphan nuclear receptor gene hzf-3/nurr1 during spatial discrimination learning. Neurobiol Learn Mem 74, 161–178 (2000). https://doi.org:10.1006/nlme.1999.3952

85 von Hertzen, L. S. & Giese, K. P. Memory reconsolidation engages only a subset of immediate-early genes induced during consolidation. J Neurosci 25, 1935–1942 (2005). https://doi.org:10.1523/JNEUROSCI.4707-04.2005

86 Ishizuka, N., Weber, J. & Amaral, D. G. Organization of intrahippocampal projections originating from CA3 pyramidal cells in the rat. J Comp Neurol 295, 580–623 (1990). https://doi.org:10.1002/cne.902950407

87 Holt, C. E. & Schuman, E. M. The central dogma decentralized: new perspectives on RNA function and local translation in neurons. Neuron 80, 648–657 (2013). https://doi.org:10.1016/j.neuron.2013.10.036

88 Tyan, S. W., Tsai, M. C., Lin, C. L., Ma, Y. L. & Lee, E. H. Serum- and glucocorticoid-inducible kinase 1 enhances zif268 expression through the mediation of SRF and CREB1 associated with spatial memory formation. J Neurochem 105, 820–832 (2008). https://doi.org:10.1111/j.1471-4159.2007.05186.x

89 Zhang, M. et al. Spatially resolved cell atlas of the mouse primary motor cortex by MERFISH. Nature 598, 137–143 (2021). https://doi.org:10.1038/s41586-021-03705-x

90 Dobin, A. & Gingeras, T. R. Mapping RNA-seq Reads with STAR. Curr Protoc Bioinformatics 51, 11 14 11–11 14 19 (2015). https://doi.org:10.1002/0471250953.bi1114s51

91 Liao, Y., Smyth, G. K. & Shi, W. featureCounts: an efficient general purpose program for assigning sequence reads to genomic features. Bioinformatics 30, 923–930 (2014). https://doi.org:10.1093/bioinformatics/btt656

92 Risso, D., Schwartz, K., Sherlock, G. & Dudoit, S. GC-content normalization for RNA-Seq data. BMC Bioinformatics 12, 480 (2011). https://doi.org:10.1186/1471-2105-12-480

93 Risso, D., Ngai, J., Speed, T. P. & Dudoit, S. Normalization of RNA-seq data using factor analysis of control genes or samples. Nat Biotechnol 32, 896–902 (2014). https://doi.org:10.1038/nbt.2931

94 Robinson, M. D., McCarthy, D. J. & Smyth, G. K. edgeR: a Bioconductor package for differential expression analysis of digital gene expression data. Bioinformatics 26, 139–140 (2010). https://doi.org:10.1093/bioinformatics/btp616

